# Antimicrobial Susceptibility Test by Surface-enhanced Raman Scattering of Bacterial Metabolites directly from Positive Blood Cultures

**DOI:** 10.1101/2022.03.23.485571

**Authors:** Yin-Yi Han, Jann-Tay Wang, Wei-Chih Cheng, Ko-Lun Chen, Yi Chi, Lee-Jene Teng, Juen-Kai Wang, Yuh-Lin Wang

## Abstract

Effective management of sepsis requires timely administration of appropriate antibiotics; therefore, a reliable and rapid antimicrobial susceptibility test (AST) is crucial. To meet clinical needs, we developed a novel AST, referred to as SERS-AST, based on the surface-enhanced Raman Scattering (SERS) technology. In this study, we applied SERS-AST to eight most common pathogens causing bacteremia, including *Staphylococcus aureus, Staphylococcus epidermidis, Enterococcus faecalis, E. faecium, Escherichia coli, Enterobacter cloacae, Klebsiella pneumoniae*, and *Acinetobacter baumannii*. Seven different antibiotics were tested, including oxacillin, levofloxacin, vancomycin, ampicillin, cefotaxime, ceftazidime, levofloxacin, and imipenem. SERS-AST determines antibiotic susceptibility of bacteria directly from positive blood cultures based on variations in bacterial SERS signals derived from secreted purines and their derivatives. The whole process could be completed within 4 hours, and the agreement rates between SERS-AST and VITEK 2 results were 96% for Gram-positive bacteria and 97% for Gram-negative bacteria.

## INTRODUCTION

Sepsis is a medical emergency with a high mortality rate, and the prevalence of sepsis and costs related to its management continue to rise [1]. It is recommended that obtaining blood cultures and lactate levels before antibiotic treatment and administration of broad-spectrum antibiotics and 30 mL/kg of crystalloid fluid for hypotension be done within 3 hours of presumptive sepsis diagnosis [2]. Timely appropriate antibiotic administration is crucial for the outcomes of patients with sepsis [3], but it relies on fast pathogen identification and antimicrobial susceptibility testing (AST). However, conventional bacterial culture and AST take 3-5 days to complete even with the aid of modern automatic microbiological analysis systems [4]. As empirical antibiotic usage may not be effective and may contribute to the development of antibiotic-resistant bacterial strains [5], there is an urgent need to develop rapid and reliable AST methods to precisely guide early antibiotic treatment for sepsis.

During the past two decades, tremendous progress has been made in the development of rapid microbiological diagnostics, such as matrix-assisted laser desorption ionization-time of flight mass spectroscopy [6], next-generation sequencing [7], nucleic acid amplification technologies [8], and single-molecule detection. These techniques can provide rapid identification and determine antibiotic susceptibility of causal organisms even for those that are non-culturable or in low concentrations [9]. However, the accuracy of these diagnostics may be impaired by unsatisfactory specimen preparations, interference of co-existing human DNA fragments or proteins, or the inability to distinguish dead from live bacteria. Furthermore, most genome-or proteome-based methods are not capable of detecting emerging antibiotic-resistant microorganisms and determining the minimum inhibitory concentration (MIC) of the antibiotics to a certain pathogen.

Surface-enhanced Raman scattering (SERS) is an emerging optical technique for rapid microbiological testing with minimal sample preparation [10]. It has evolved to become a reliable and sensitive method for qualitative and quantitative analyses of various substances. We have used it to identify various *Mycobacterium* species [11], determine antibiotic susceptibility of certain bacteria [12, 13], and monitor environmental pollution [14] and food safety [15].

The SERS spectra of bacteria determined in our studies [12, 13, 16] are similar to those obtained by other groups [17-19]. Bacterial SERS signals are derived from secreted purines and their derivatives (e.g., adenine, hypoxanthine, xanthine, guanine, uric acid, and adenosine monophosphate) [20, 21]. We used SERS signal patterns, such as the 730-cm^-1^ peak for *Staphylococcus aureus* and the 724-cm^-1^ peak for *Escherichia coli*, to determine changes in the concentration of live bacteria after an antibiotic treatment [12]. The consistency of our SERS-AST results on *S. aureus* and *E. coli* isolated from blood cultures was 93% compared with those of conventional AST methods [13].

To apply our SERS-AST to bacteria causing sepsis, more tests on various bacterium-antibiotic combinations are required. As SERS-AST is based on variations in bacterial metabolism in response to antibiotic treatment [22, 23], protocol modifications are needed due to different metabolic activities of various bacteria and mechanisms of action of various antibiotics. In this study, we developed SERS-AST protocols to examine the eight most common bacteria causing sepsis [24, 25] for their responses to commonly used antibiotics [26].

## MATERIALS AND METHODS

### Study Design

This study was aimed to improve our previous SERS-AST method for determination of antibiotic susceptibility of bacteria causing bloodstream infections [13], including *Staphylococcus aureus, Staphylococcus epidermidis, Enterococcus faecalis, Enterococcus faecium, E. coli, Enterobacter cloacae, Klebsiella pneumoniae*, and *Acinetobacter baumannii*. Seven antibiotics were tested, including oxacillin (OXA), levofloxacin (LVX), vancomycin (VAN), ampicillin (AMP), cefotaxime (CTX), ceftazidime (CAZ), levofloxacin (LVX), and imipenem (IMP). SERS-AST results were verified by comparing them with those determined by the VITEK 2 automatic microbial diagnostic system (bioMérieux, Hazelwood, Missouri, USA).

### Ethical Approval Statement

This study was conducted according to the guidelines of the Declaration of Helsinki-based Ethical Principles for Medical Research and was approved by the Ethics Committee of National Taiwan University Hospital (NTUH) (approval number: 201107031RC). Written informed consent was obtained from each patient before the study was initiated. Patient confidentiality was strictly protected.

### Preparation of blood samples

SERS-AST was performed as described previously [13]. The entire test included blood sample preparation, antibiotic treatment, SERS measurement, and receiver operating characteristic (ROC) analysis. Five ml of a positive blood culture was used for each SERS-AST. To prevent blood components, such as hemoglobin [27], from interfering with SERS measurements, red blood cells were selectively lysed with Ammonium-Chloride-Potassium (ACK) buffer [28] followed by ultrasonication to facilitate sonoporation [29]. The bacteria in the ACK-treated sample were pelleted, resuspended in Mueller-Hinton broth (MHB) to 2.5 × 10^7^ CFU/ml, treated with or without a selected antibiotic at various concentrations, and incubated at 37°C until the OD_600_ value of the untreated control culture reached 1 or greater; this usually took 2 to 3 hours. The bacteria were then pelleted and resuspended in deionized water to 3.0 × 10^9^ CFU/ml for the non-antibiotic-treated control sample prior to SERS-AST. Bacterial cells in each antibiotic-treated sample were resuspended with the same amount of deionized water as the control sample. For *A. baumannii*, the cell suspension in deionized water was further incubated in a shaking water bath at 37°C for 30 minutes to ensure sufficient amounts of secreted purine derivatives for SERS-AST.

### Antibiotic treatment

Various antibiotic concentrations were tested as follows: 0.0625, 0.125, 0.25, and 0.5 μg/ml for the *S. epidermidis*-OXA combination; 0.5, 1, 2, and 4 μg/ml for *S. aureus-*OXA, *S. aureus*-LVX, *S. epidermidis*-LVX, *E. coli*-CTX, and *E*. cloacae-CTX combinations; 1, 2, 4, and 8 μg/ml for *E. faecalis*-LVX, *E. faecium*-LVX, *E. coli*-LVX, *K. pneumoniae*-CTX, *K. pneumoniae*-IPM, *A. baumannii*-LVX, *A. baumannii*-IPM combinations; 2, 4, 8, and 16 μg/ml for *E. faecium*-VAN, *E. coli*-CAZ, *E. cloacae*-CAZ, *K. pneumoniae*-CAZ, *K. pneumoniae*-LVX combinations; and 4, 8, 16, and 32 μg/ml for *E. faecium*-AMP and *A. baumannii*-CAZ combinations.

### SERS measurement and spectral analysis

The SERS device was a glass slide coated with anodic aluminum oxide with embedded Ag nanoparticles (AgNP/AAO) [30]. For SERS assay, the AgNP/AAO slide was placed inside an aluminum trough, which holds the slide in place. A 1-mm thick aluminum sheet with 2 rows of 4 holes of 1.5-mm in diameter separated from each other by 2.5 mm was hung on the edges of the trough covering the slide 7 mm above. One microliter of each bacterial suspension was then placed onto the slide with a pipet to deliver it through the holes on the aluminum sheet with the 4 antibiotic-treated samples forming 4 spots in the top row on the slide and the 4 non-treated control samples forming another 4 spots in the bottom row at precise locations. The slide was then removed from the trough and placed on a hot plate to dry the samples at 55°C for 15 minutes before SERS measurements.

The SERS instrument was composed of a He-Ne laser emitting a 632.8 nm light, an upright optical microscope (BX61WI, Olympus), a Raman probe (Superhead, Horiba), a spectrometer (HE 633, Horiba), and a thermoelectric-cooled charge-coupled (CCD) camera. The laser beam was delivered through an optical fiber into a 20× objective lens by the Raman probe. The scattered light that bounced back from the sample on the AgNP/AAO slide was collected and delivered by the same objective lens through another optical fiber to the spectrometer and CCD for spectral recording and analyses.

The height of the dominant peak in a SERS spectrum was used to determine antibiotic susceptibility. For Gram-positive bacteria, a dominant spectral peak at 730 cm^−1^ was observed due to secreted adenine. For Gram-negative bacteria, a dominant spectral peak at 724 cm^−1^ was observed because of secreted hypoxanthine. For *A. baumannii*, which secrets primarily xanthine [20], a dominant spectral peak at 654 cm^-1^ was observed. The SERS-AST signal ratio was calculated by dividing the peak signal height value of the antibiotic-treated sample by that of the adjacent non-treated control. For each bacterial-antibiotic test, four SERS-AST signal ratios corresponding to four different antibiotic concentrations were obtained and used to perform ROC analyses.

### Receiver operating characteristic analysis

ROC analysis was performed to determine the optimal cutoff signal ratio for antibiotic susceptibility of the bacterium in each bacterium-antibiotic combination. The optimal cutoff signal ratio (*r*_*op*_) was the ratio that maximized the Youden index (J = sensitivity + specificity −1) and accounted for all antibiotic-resistant samples. As ROC is a binary classification system (i.e., resistant or susceptible), a bacterium determined to be intermediate susceptible to an antibiotic was classified as resistant. For a bacterium-antibiotic test, the antibiotic concentration that yielded a signal ratio smaller than *r*_*op*_ indicated that the bacteria in the sample were inhibited at that concentration. The lowest antibiotic concentration at which the inhibition occurred was defined as the MIC of the tested bacterium. The final antibiotic susceptibility of the tested bacterium was determined by comparing the MIC to that of the AST standards of the Clinical and Laboratory Standards Institute (CLSI) [31].

### Comparative evaluation of SERS-AST results with clinical standards

The results of SERS-AST were compared with those determined by the automatic VITEK 2 system and were classified as agreement and disagreement.

## RESULTS

From March 2016 to June 2019, a total of 164 bacterial isolates were analyzed from blood samples, including *S. aureus* (n=20), *S. epidermidis* (n=21), *E. faecalis* (n=20), *E. faecium* (n=21), *E. coli* (n=20), *E. cloacae* (n=20), *K. pneumoniae* (n=21) and *A. baumannii* (n=21). Three samples failed to generate analyzable SERS signals, including one *S. epidermidis* and one *E. faecium* samples with insufficient growth and one *A. baumannii* sample that generated unrecognizable SERS signals. Results of three *K. pneumoniae*-IPM tests were also excluded because of improper antibiotic preparation. There were 141 bacterium-antibiotic tests excluded from ROC analyses due to insufficient numbers (< 3) of antibiotic-resistant samples, including *S. aureus*-VAN, *S. epidermidis*-VAN, *E. faecalis*-AMP, *E. faecalis-*VAN, *E. coli*-IPM, *E. cloacae*-LVX, and *E. cloacae*-IPM tests. A total of 401 datasets were analyzed by ROC.

### SERS-AST for Gram-positive Bacteria

The SERS-AST signal ratio (signal value of treated sample divided by that of untreated) of spectral peaks at 730 cm^-1^ (*r*_730_) was used to analyze antibiotic response of each Gram-positive bacterium (Figure 1). For *S. aureus* treated with OXA or LVX, the patterns of *r*_730_ value variation were similar for susceptible isolates with a low *r*_730_ value at the susceptible breakpoint and higher concentrations, while the *r*_730_ values of resistant isolates remained high for all 4 antibiotic concentrations. The areas under the ROC curve (AUC) were 0.99 for OXA and 0.95 for LVX (Figure 3). There was one disagreement in each of *S. aureus*-OXA and *S. aureus*-LVX tests, in which SERS-AST determined the isolate as resistant, but VITEK 2 determined it as susceptible. The agreement rate between the results of SERS-AST and VITEK 2 was 95% for both tests (Table 1). Good performance of SERS-AST was also observed in *S. epidermidis* treated with OXA or LVX and *E. faecium* treated with AMP or VAN. Only one disagreement was seen in *S. epidermidis*-LVX tests, in which SERS-AST determined the isolate as resistant, whereas VITEK 2 determined it as susceptible. The agreement rate between the results of SERS-AST and VITEK 2 was 100% for *S. epidermidis*-OXA, *E. faecium*-AMP, and *E. faecium*-VAN tests, and 95% for *S. epidermidis*-LVX tests. The SERS-AST results of *E. faecalis* and *E. faecium* treated with LVX were less satisfactory as low AUC values (0.90 for *E. faecalis*-LVX tests and 0.67 for *E. faecium*-LVX tests) and low agreement rates with the results of VITEK 2 (85% for *E. faecalis*-LVX tests and 95% for *E. faecium*-LVX tests) were observed. There were 3 disagreements for *E. faecalis*-LVX tests and one disagreement for *E. faecium*-LVX tests, in which SERS-AST determined the isolates as resistant, while VITEK 2 determined them as susceptible. The overall agreement rate between SERS-AST and VITEK-2 results of the four Gram-positive bacteria was 96%.

**Table 1.** SERS-AST results of Gram-positive bacteria.

| Bacterial species | Antibiotic | Number of S and R<br>isolates determined<br>by VITEK 2 | | $r_{op}$ | AUC<br>value | Disagreement<br>(No.) | Agreement rate<br>(%) |
| --- | --- | --- | --- | --- | --- | --- | --- |
|  |  | S | R |  |  |  |  |
| *S. aureus* | OXA | 12 | 8 | 0.36 | 0.99 | 1 | 95 |
|  | LVX | 16 | 4 | 0.83 | 0.95 | 1 | 95 |
| *S. epidermidis* | OXA | 6 | 14 | 0.25 | 1 | 0 | 100 |
|  | LVX | 12 | 8 | 0.43 | 0.99 | 1 | 95 |
| *E. faecalis* | LVX | 17 | 3 | 0.68 | 0.90 | 3 | 85 |
| *E. faecium* | LVX | 3 | 17 | 0.61 | 0.67 | 1 | 95 |
|  | VAN | 4 | 16 | 0.33 | 1 | 0 | 100 |
|  | AMP | 9 | 11 | 0.44 | 1 | 0 | 100 |

S and R stand for susceptible and resistant, respectively. $r_{op}$ is the optimal cut-off SERS-AST signal ratio. The bacteria treated with a certain antibiotic concentration that yielded a SERS-AST signal ratio lower than $r_{op}$ were considered as being inhibited at that concentration. AUC is the area under the receiver operating characteristic (ROC) curve. Numbers of isolates with disagreement results and agreement rate (%) were determined by comparing the results of SERS-AST with those of VITEK 2.

**Figure 1.**
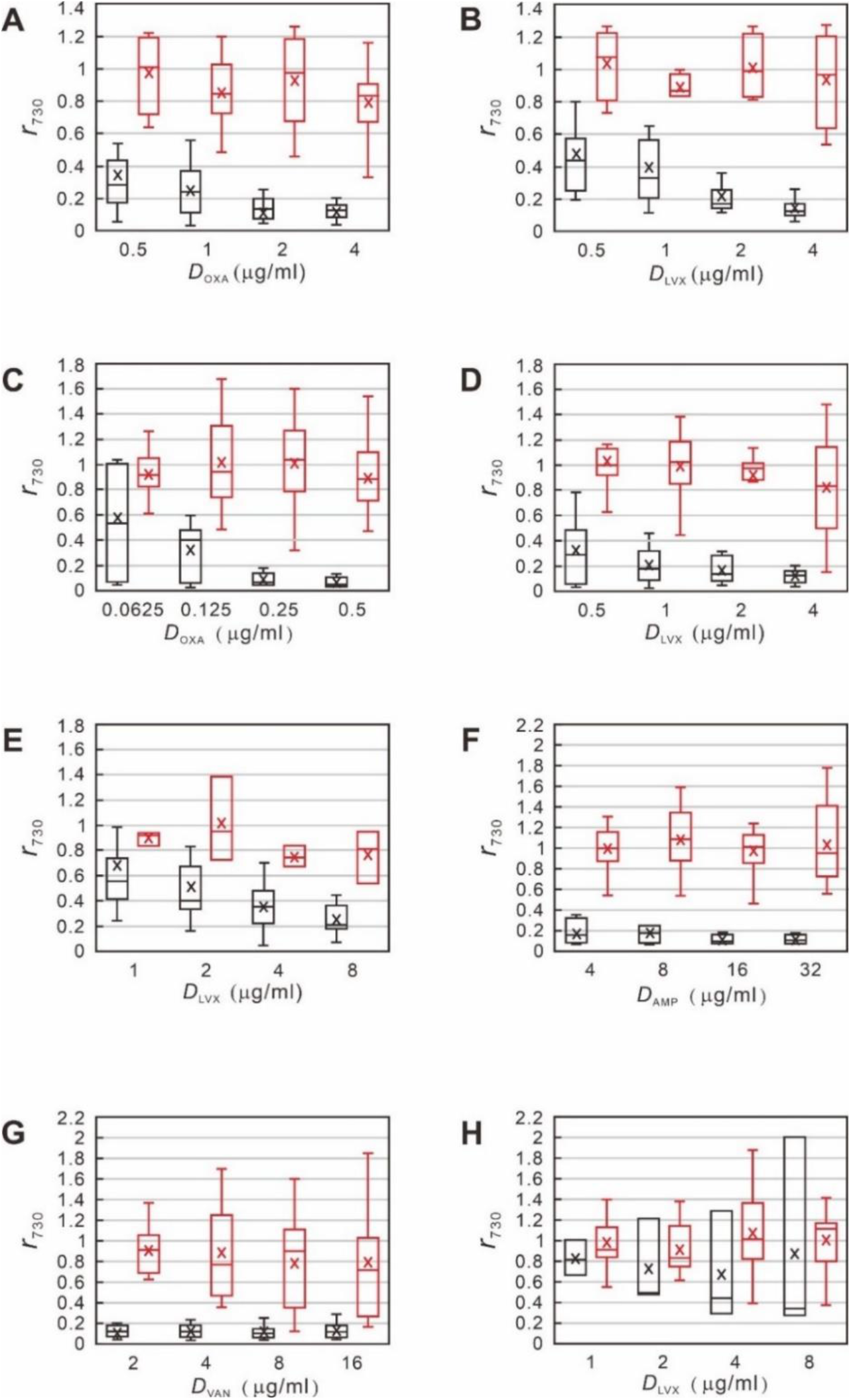
Box-and-whisker plots of SERS-AST results of Gram-positive bacteria treated with various concentration of antibiotics. **A**. *S. aureus* with OXA, **B**. *S. aureus* with LVX, **C**. *S. epidermidis* with OXA, **D**. *S. epidermidis* with LVX, **E**. *E. faecalis* with LVX, **F**. *E. faecium* with AMP, **G**. *E. faecium* with VAN, and **H**. *E. faecium* with LVX. Values on the Y-axis are SERS-AST signal ratios (***r***_**730**_) calculated by dividing the peak height value at 730 cm^-1^ of the antibiotic-treated sample by that of the non-treated control sample. Values on the X-axis are antibiotic concentrations. The boxes represent 25th to 75th percentiles of samples, with the 50th percentile indicated with a small line in the boxes. The 10th and 90th percentiles of samples are indicated with whiskers. The x letter inside each box represents the mean ***r***_**730**_ value. Red and black boxes represent the data of resistant and susceptible samples, respectively. VAN, vancomycin; OXA, oxacillin; AMP, ampicillin; LVX, levofloxacin.

For Gram-positive bacteria, the decrease in SERS-AST signal ratio as a result of LVX treatment was generally smaller than those of other antibiotic treatments (Figure 1). Therefore, a higher *r*_*op*_ for AUC was obtained (0.83 vs. 0.36 in *S. aureus*-LVX and *S. aureus*-OXA tests; 0.43 vs. 0.25 in *S. epidermidis*-LVX and *S. epidermidis*-OXA tests; 0.68 in *E. faecium-*LVX, 0.61 vs. 0.33 and 0.44 in *E. faecium*-LVX, *E. faecium*-VAN, and *E. faecium*-AMP tests) (Table 1).

### SERS-AST for Gram-negative Bacteria

The SERS-AST signal ratio (signal value of treated sample divided by that of untreated) of a spectral peak at 654 cm^-1^ (*r*_654_) was used to determine antibiotic response of *A. baumannii* and that at 724 cm^-1^ (*r*_724_) was used to analyze other Gram-negative bacteria (Figure 2). For *E. coli* treated with CTX, CAZ, or LVX, the patterns of *r*_724_ value variation were similar for susceptible isolates with a low *r*_724_ value at the susceptible breakpoint and higher concentrations, while the *r*_724_ values of resistant isolates remained high for all 4 antibiotic concentrations. The AUC values of the three antibiotic tests (CTX, CAZ, and LVX) were all 1, indicating a 100% agreement between the results of SERS-AST and VITEK 2 (Table 2). The SERS-AST was also successfully applied to other Gram-negative bacteria with an AUC value of 1 for all tests, including *K. pneumoniae*-IPM, *A. baumannii*-CAZ, *A. baumannii*-LVX, *and A. baumannii*-IPM tests. The AUC values of *E. cloacae*-CTX, *E. cloacae*-CAZ, *K. pneumoniae*-CTX, and *K. pneumoniae*-LVX tests were 0.96, 0.98, 0.98, and 0.96, respectively. There was only one disagreement in each of the 4 tests, in which SERS-AST determined the isolates as resistant, but VITEK 2 determined them as susceptible. For *K. pneumoniae*-CAZ tests, the AUC was 0.91, and there were 3 disagreements, in which SERS-AST determined the isolates as resistant, while VITEK 2 determined them as susceptible. Overall, the agreement rate between SERS-AST and VITEK 2 results was 97%.

**Table 2.** SERS-AST results of Gram-negative bacteria.

| Bacterial species | Antibiotic | Number of S and R<br>isolates determined<br>by VITEK 2 | | $r_{op}$ | AUC<br>value | Disagreement<br>(No.) | Agreement<br>rate (%) |
| --- | --- | --- | --- | --- | --- | --- | --- |
|  |  | S | R |  |  |  |  |
| *E. coli* | CTX | 14 | 6 | 0.65 | 1 | 0 | 100 |
|  | CAZ | 14 | 6 | 0.53 | 1 | 0 | 100 |
|  | LVX | 14 | 6 | 0.72 | 1 | 0 | 100 |
| *E. cloacae* | CTX | 12 | 8 | 0.68 | 0.96 | 1 | 95 |
|  | CAZ | 12 | 8 | 0.69 | 0.98 | 1 | 95 |
| *K. pneumoniae* | CTX | 12 | 9 | 0.18 | 0.98 | 1 | 95 |
|  | CAZ | 12 | 9 | 0.13 | 0.91 | 3 | 86 |
|  | LVX | 16 | 5 | 0.68 | 0.96 | 1 | 95 |
|  | IPM | 15 | 3 | 0.37 | 1 | 0 | 100 |
| *A. baumannii* | CAZ | 10 | 10 | 0.56 | 1 | 0 | 100 |
|  | LVX | 10 | 10 | 0.52 | 1 | 0 | 100 |
|  | IPM | 10 | 10 | 0.49 | 1 | 0 | 100 |

S and R stand for susceptible and resistant, respectively. $r_{op}$ is the optimal cut-off SERS-AST signal ratio. The bacteria treated with a certain antibiotic concentration that yielded a SERS-AST signal ratio lower than $r_{op}$ were considered as being inhibited at that concentration. AUC is the area under the receiver operating characteristic (ROC) curve. Numbers of isolates with disagreement results and agreement rate (%) were determined by comparing the results of SERS-AST with those of VITEK 2.

**Figure 2.**
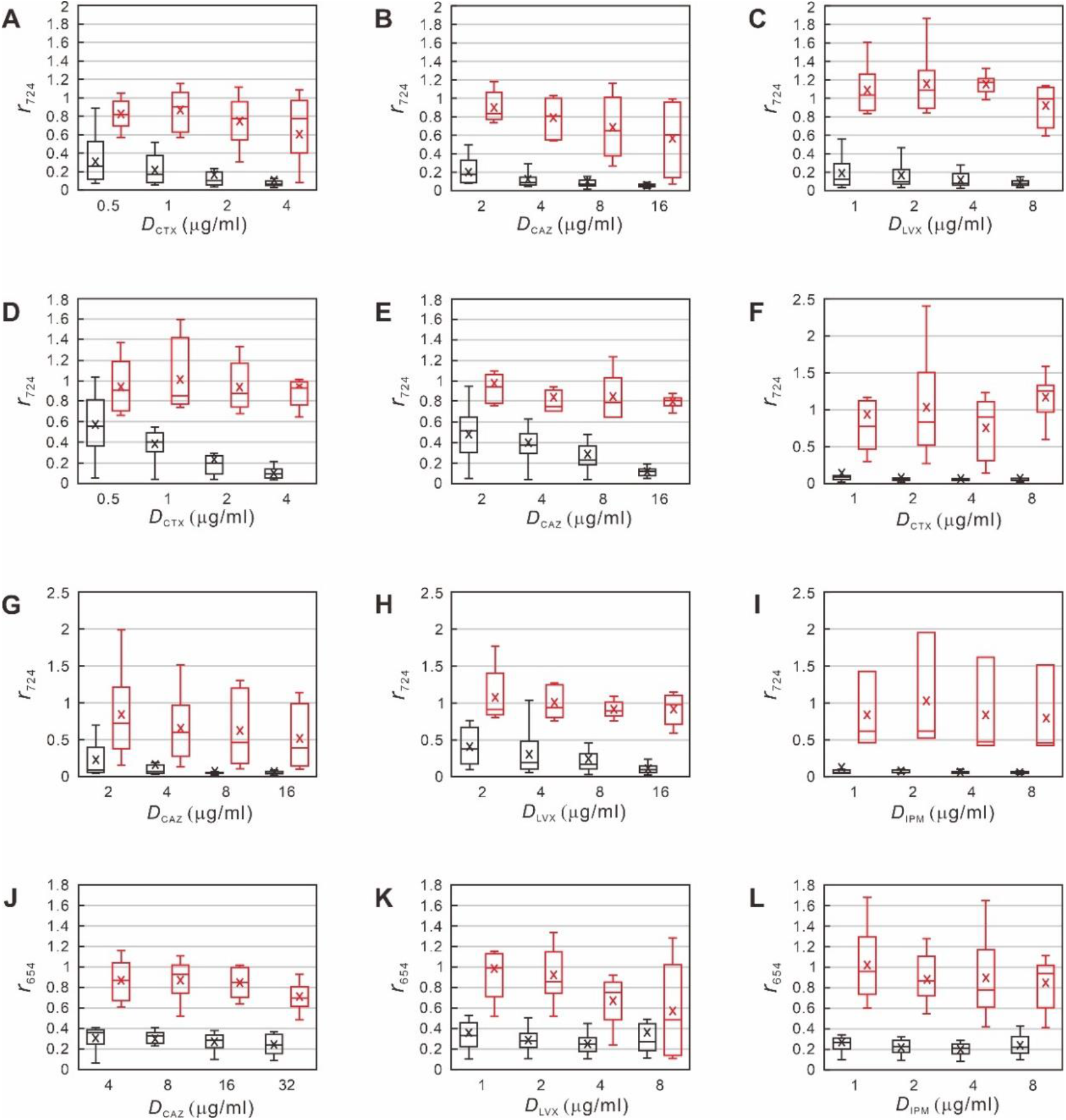
Box-and-whisker plots of SERS signals of Gram-negative bacteria treated with various concentrations of antibiotics. **A**. *E. coli* with CTX, **B**. *E. coli* with CAZ, **C**. *E. coli* with LVX, **D**. *E*. cloacae with CTX, **E**. *E. cloacae* with CAZ, **F**. *K. pneumoniae* with CTX, **G**. *K. pneumoniae* with CAZ, **H**. *K. pneumoniae* with LVX, **I**. *K. pneumoniae* with IPM, **J**. *A. baumannii* with CAZ, **K**. *A. baumannii* with LVX, **L**. *A. baumannii* with IPM. For **A**-**I**, values on the Y-axis are SERS-AST signal ratios (***r***_**724**_) calculated by dividing the peak height value at 724 cm^-1^ of the antibiotic-treated sample by that of the non-treated control sample. For **J**-**L**, values on the Y-axis are SERS-AST signal ratios (***r***_**654**_) calculated by dividing the peak height value at 654 cm^-1^ of the antibiotic-treated sample by that of the non-treated control sample. Values on the X-axis are antibiotic concentrations. The boxes represent 25th to 75th percentiles of samples, with the 50th percentile indicated with a small line in the boxes. The 10th and 90th percentiles of samples are indicated with whiskers. The x letter inside each box represents the mean ***r***_**724**_ or ***r***_**654**_ value. Red and black boxes represent the data of resistant and susceptible samples, respectively. LVX, levofloxacin, CTX, cefotaxime, CAZ, ceftazidime, and IPM, imipenem.

**Figure 3.**
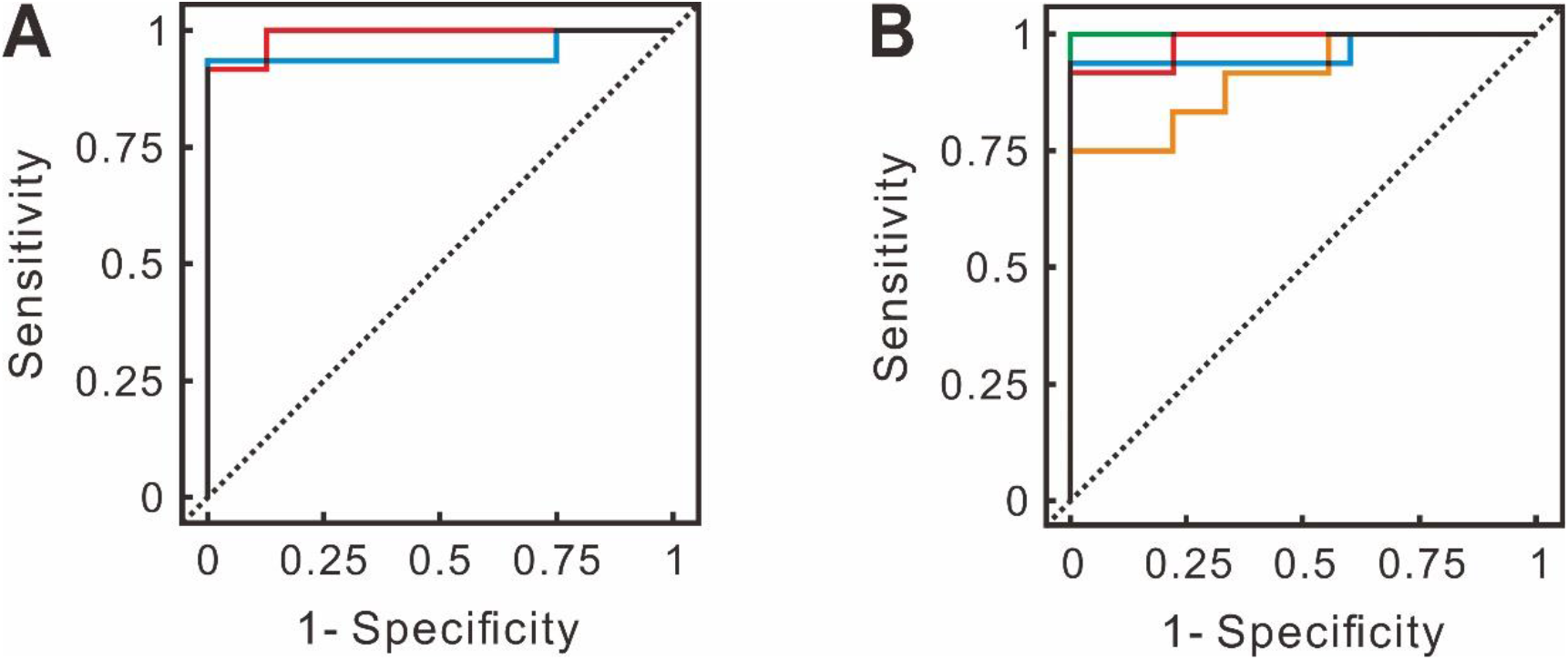
Representative ROC curves of SERS-AST results. **A**. Results of *S. aureus* treated with OXA or LVX. Red and blue curves represent results of OXA and LVX treatments, respectively. The black segments indicate overlaps between the two results. **B**. Results of *K. pneumoniae* treated with CTX, LVX, IPM, or CAZ. Red, blue, green, and orange curves represent results of CTX, LVX, IPM, and CAZ treatments, respectively. The black segments indicate overlaps among the results.

## DISCUSSION

In this study, we extended our previous proof-of-principle study of SERS-AST, which was performed on *S. aureus*-OXA and *E. coli*-CTX combinations [13], to 20 bacterium-antibiotic combinations of eight most common pathogens causing bacteremia and seven commonly used antibiotics. The whole SERS-AST process could be completed within 4 hours, and the agreement rates between results of SERS-AST and VITEK 2 were 96% for Gram-positive bacteria and 97% for Gram-negative bacteria. The time needed for a SERS-AST is significantly shorter than that for VITEK 2, which requires subculture of a positive blood culture sample on agar plates to obtain pure organisms. In contrast, SER-AST can directly assay positive blood cultures without additional cultures.

In SERS-AST, bacterial response to antibiotics is determined by changes in SERS peak height, while conventional ASTs are mainly based on changes in the density (OD_600_ value) of antibiotic-treated cultures. In this study, we found that approximately 30% of the results determined by OD_600_ values of antibiotic-treated bacterial cultures did not agree with those of SERS-AST or VITEK 2. Most of the disagreements were of Gram-negative bacteria treated with *β*-lactam antibiotics, such as *E. cloacae*-CTX tests (Figure 4). It has been shown that an initial response of *E. cloacae* to treatment with a *β*-lactam antibiotic is elongation of cells, instead of decreased number of cells [32]. Therefore, the OD_600_ value of the treated culture may not change within 2 hours of treatment. Such cell morphological change is postulated to be a repair process for survival [33]. In contrast, SERS-AST measures the amounts of secreted purines and their derivatives in response to an antibiotic treatment. Such measurements are less affected by changes in cell morphology as in the case of our *E. cloacae*-CTX tests.

**Figure 4.**
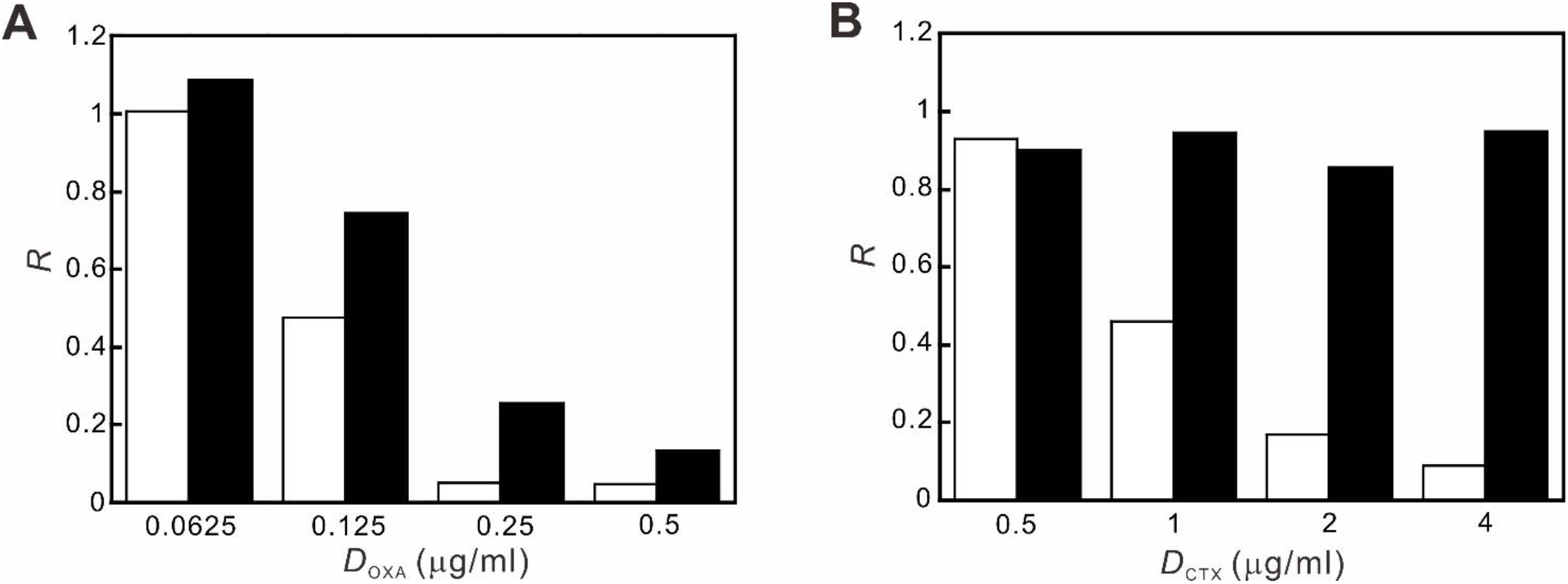
SERS-AST signals versus optical density of antibiotic-treated cultures. **A**. Variations in SERS-AST signal ratios and OD_600_ ratios of OXA-susceptible *S. epidermidis* cultures. **B**. Variations in SERS signal ratios and OD_600_ ratios CTX-susceptible *E. cloacae* cultures. Values on the Y-axis are ratios of OD_600_ or SERS peak height values calculated by dividing the value of the antibiotic-treated sample by that of the non-treated control sample. Values on the X-axis are antibiotic concentrations.

Several modifications of the SERS-AST protocol were done to obtain reliable results from certain bacteria, e.g., *A. baumannii*, which failed to generate recognizable SERS signals even when the incubation time of antibiotic treatments was extended to 3 hours. This problem may be due to the low permeability (approximately 1/100 that of *E. coli*) of its outer membrane to small molecules, thus hindering the secretion of xanthine [34, 35]. To determine the optimal condition for *A. baumannii* SERS-AST, the antibiotic-treated cell suspension in water was incubated in a shaking water bath at 25, 37, or 50°C for 30, 60, or 90 minutes. As the result showed that an additional 30-minute incubation in a shaking water bath at 37°C rendered a 2.2-fold increase in SERS signal, this condition was used for all subsequent SERS-AST tests for *A. baumannii*. We postulate that this signal improvement is mainly due to the impact of placing bacteria in a nutrient-deficient environment as this starvation process has been shown to stimulate more secretion of purines and their derivatives [36-38].

In this study, 14 SERS-AST tests gave results that did not agree with those of VITEK 2, including 3 each of *E. faecalis*-LVX and *K. pneumoniae*-CAZ tests, and one each of *S. aureus*-OXA, *S. aureus*-LVX, *S. epidermidis*-LVX, *E. faecium*-LVX, *E. cloacae*-CTX, *E. cloacae-*CAZ, *K. pneumoniae*-CTX, and *K. pneumoniae*-LVX tests (Table 1, 2). Seven (50%) of the 14 cases were of LVX tests with mostly Gram-positive bacteria (Table 1). Among the seven antibiotics tested, LVX is the only one not acting on cell wall synthesis. It is a quinolone antibiotic that inhibits gyrase and topoisomerase IV leading to impaired DNA replication, repair, and recombination [39]. It has been shown that gyrase is the primary target of quinolones in Gram-negative bacteria, while topoisomerase IV is the main target of quinolones in Gram-positive bacteria [39]. In DNA replication, inhibition occurs within minutes when the antibiotic acts on gyrase [40] but takes place later if it targets topoisomerase IV [41]. However, bacterial responses to quinolones have been shown to vary on a species-by-species and drug-by-drug basis [42]. It is likely that a longer time is required for LVX to inhibit the growth of slow-growing Gram-positive bacteria, such as *E. faecalis*. In an attempt to optimize the condition for *E. faecium-*LVX tests, we obtained more clear-cut SERS-AST results when the antibiotic treatment time was extended to 3 or 4 hours. Based on these results, we recommend that the antibiotic treatment time be extended to 3 hours for all LVX tests in future studies

Five (36%) of the 14 tests with SERS-AST results not consistent with the results of VITEK 2 involved *K. pneumoniae* with antibiotics CAZ, CTX, and LVX (Table 2). *K. pneumoniae* is known to produce a pronounced polysaccharide capsule covering the entire bacterial surface resulting in a mucoid phenotype [43] with reduced ability to secrete purines and their derivatives, thus yielding weak SERS signals. *K. pneumoniae* is also known to produces CTX M β-lactamase, which is an extended-spectrum β-lactamase (ESBL) and can degrade third-generation cephalosporin antibiotics at different rates [44]. Compared to CTX, which is the preferred target of CTX M β-lactamase, CAZ is relatively resistant to that enzyme and may require more than 2 hours to be degraded. Therefore, some CAZ-resistant bacteria may be determined by laboratory testing as susceptible, which is inconsistent with clinical manifestations. The Advanced Expert System (AES) of VITEK 2 can identify bacteria with ESBL by special software and modify the primary laboratory results accordingly [45]. In this study, there were five AES-revised *K. pneumoniae*-CAZ results. Three of such cases that were determined as susceptible by SERS-AST after the 2-hour antibiotic treatment were interpreted by AES as resistant.

This study has several strengths. It demonstrated the possibility of performing SERS-AST on clinically significant bacteria directly from positive blood cultures. This study also successfully developed methods, such as starvation, to perform SERS-AST on bacteria (i.e., *A. baumannii*) that were previously not assayable. The finding that extending the LVX treatment time to 3 hours, especially for Gram-positive bacteria such as *E. faecalis* that grows slowly, can generate satisfactory SERS-AST results is very significant.

To move the SERS-AST forward for clinical applications, further improvements are needed. Special methods for processing samples with multiple organisms and those with other microorganisms such as *Pseudomonas aeruginosa* and fungi remain to be developed. Although *P. aeruginosa* is also a major bacterium causing sepsis, we have not been able to perform SERS-AST on it because of interference by its fluorescent pigments. As SERS-AST is based on changes in bacterial metabolism due to antibiotic treatment, it is possible to modulate the metabolic activity of bacteria by altering their growth environments with substances such as culture media and cations (e.g., Mg^2+^, Ca^2+^ and Na^+^) [46] or with pH adjustment [33]. The SERS-AST instrument that we have used is a prototype with many rooms for improvements to increase it sensitivity and specificity.

## CONCLUSIONS

In this study, we performed SERS-based rapid ASTs on 20 bacterium-antibiotic combinations of eight most common pathogens causing bacteremia and seven commonly used antibiotics. Distinct from culture-based and genome-or proteome-based ASTs, SERS-AST analyzes changes in bacterial metabolism due to antibiotic treatment by measuring the amounts of secreted purines and their derivative. The entire SERS-AST includes the following steps in sequence: obtain 5 ml aliquot of a positive blood culture after completion of bacterial identification, lyse red blood cells in the sample, wash bacteria and resuspend them in MHB with various concentrations of the antibiotic to be tested, incubate the mixtures at 37°C for 2 - 3 hours, pellet bacterial cells and resuspend them in deionized water, examine the samples by SERS, and then analyze the obtained SERS signal ratio by ROC. The whole process can be completed within 4 hours. The agreement rates between our SERS-AST results and VITEK 2 results were 96% for Gram-positive bacteria and 97% for Gram-negative bacteria.

## Acknowledgements

We thank the following individuals in various departments of National Taiwan University Hospital: Dr. Chin-Hao Chang in the Center of Statistical Consultation and Research, Department of Medical Research for statistical consultation; Dr. Shu-Yi Huang in the Department of Medical Research for suggestions in manuscript preparation; and staffs of the Department of Laboratory Medicine for providing samples and VITEK 2 results. We also thank the Seventh Core Laboratory and Center for Infection Control for providing Laboratory facility.

## Author contributions

Y-Y. H., J-K. W., and Y-L.W. designed and conducted the study. W-C. C., K-L. C., and Y. C. performed experiments. Y-Y. H. and J-K. W. interpreted results and wrote the manuscript. L-J.T. and C-T. W. participated in the analysis of results. Y-L.W. reviewed results and edited the manuscript.

## Funding

This study was supported by grants from the Ministry of Science and Technology (MOST 106-2745-M-001-004 -ASP, MOST 107-2745-M-001-004 -ASP, MOST 108-2639-M-001-003 -ASP, MOST 109-2639-M-001-005 -ASP).

## Competing interests

The authors declare no competing interests.

## REFERENCES

1. Goto, M. and M.N. Al-Hasan, Overall burden of bloodstream infection and nosocomial bloodstream infection in North America and Europe. Clin Microbiol Infect, 2013. 19(6): p. 501–9.

2. Seymour, C.W., et al., Time to Treatment and Mortality during Mandated Emergency Care for Sepsis. N Engl J Med, 2017. 376(23): p. 2235–2244.

3. Retamar, P., et al., Impact of inadequate empirical therapy on the mortality of patients with bloodstream infections: a propensity score-based analysis. Antimicrob Agents Chemother, 2012. 56(1): p. 472–8.

4. Webb, B.J., B. Jones, and N.C. Dean, Empiric antibiotic selection and risk prediction of drug-resistant pathogens in community-onset pneumonia. Curr Opin Infect Dis, 2016. 29(2): p. 167–77.

5. Eliopoulos, G.M., D.L. Paterson, and L.B. Rice, Empirical Antibiotic Choice for the Seriously Ill Patient: Are Minimization of Selection of Resistant Organisms and Maximization of Individual Outcome Mutually Exclusive? Clinical Infectious Diseases, 2003. 36(8): p. 1006–1012.

6. Oviano, M. and G. Bou, Matrix-Assisted Laser Desorption Ionization-Time of Flight Mass Spectrometry for the Rapid Detection of Antimicrobial Resistance Mechanisms and Beyond. Clin Microbiol Rev, 2019. 32(1).

7. van Belkum, A. and W.M. Dunne, Jr., Next-generation antimicrobial susceptibility testing. J Clin Microbiol, 2013. 51(7): p. 2018–24.

8. Lee, S.H., et al., Emerging ultrafast nucleic acid amplification technologies for next-generation molecular diagnostics. Biosensors and Bioelectronics, 2019. 141: p. 111448.

9. Riedel, S. and K.C. Carroll, Early Identification and Treatment of Pathogens in Sepsis: Molecular Diagnostics and Antibiotic Choice. Clin Chest Med, 2016. 37(2): p. 191–207.

10. Jarvis, R.M. and R. Goodacre, Discrimination of bacteria using surface-enhanced Raman spectroscopy. Anal. Chem., 2004. 76: p. 40–47.

11. Cheng, W.-C., et al., Sensible Functional Linear Discriminant Analysis Effectively Discriminates Enhanced Raman Spectra of Mycobacterium Species. Analytical Chemistry, 2021. 93(5): p. 2785–2792.

12. Liu, C.Y., et al., Rapid bacterial antibiotic susceptibility test based on simple surface-enhanced Raman spectroscopic biomarkers. Sci Rep, 2016. 6: p. 23375.

13. Han, Y.-Y., et al., Rapid antibiotic susceptibility testing of bacteria from patients’ blood via assaying bacterial metabolic response with surface-enhanced Raman spectroscopy. Scientific Reports, 2020. 10(1): p. 12538.

14. Dvoynenko, O., et al., Speciation Analysis of Cr(VI) and Cr(III) in Water with Surface-Enhanced Raman Spectroscopy. ACS Omega, 2021. 6(3): p. 2052–2059.

15. Lian, W.N., et al., Rapid detection of copper chlorophyll in vegetable oils based on surface-enhanced Raman spectroscopy. Food Addit Contam Part A Chem Anal Control Expo Risk Assess, 2015. 32(5): p. 627–34.

16. Liu, T.T., et al., A high speed detection platform based on surface-enhanced Raman scattering for monitoring antibiotic-induced chemical changes in bacteria cell wall. hPLoS One, 2009. 4(5): p. e5470.

17. Premasiri, W.R., et al., Characterization of the surface enhanced raman scattering (SERS) of bacteria. J Phys Chem B, 2005. 109(1): p. 312–20.

18. Boardman, A.K., et al., Rapid Detection of Bacteria from Blood with Surface-Enhanced Raman Spectroscopy. Anal Chem, 2016. 88(16): p. 8026–35.

19. Zhao, X., M. Li, and Z. Xu, Detection of Foodborne Pathogens by Surface Enhanced Raman Spectroscopy. Frontiers in microbiology, 2018. 9: p. 1236–1236.

20. Premasiri, W.R., et al., The biochemical origins of the surface-enhanced Raman spectra of bacteria: a metabolomics profiling by SERS. Anal Bioanal Chem, 2016. 408(17): p. 4631–47.

21. Chiu, S.W.Y. and C.Z. Cheng HW, Wang HH, Lai MY, Wang JK, Wang YL, Quantification of biomolecules responsible for biomarkers in the surface-enhanced Raman spectra of bacteria using liquid chromatography-mass spectrometry. Phys. Chem. Chem. Phys, 2018. 20: p. 8032–8041

22. Belenky, P. and P.C. Ye Jd, Cohen NR, Lobritz MA, Ferrante T, Jain S, Korry BJ, Schwarz EG, Walker GC, Collins JJ., Bactericidal antibiotics induce toxic metabolic perturbations that lead to cellular damage. Cell Rep., 2015. 13: p. 968–980

23. Zampieri, M., et al., Nontargeted Metabolomics Reveals the Multilevel Response to Antibiotic Perturbations. Cell Rep, 2017. 19(6): p. 1214–1228.

24. Infection Control Center, N.T.U.H.N., Annual Report of Nosocomial Infection. 2015: NTUH, Taipei.

25. Diekema, D.J., et al., The Microbiology of Bloodstream Infection: 20-Year Trends from the SENTRY Antimicrobial Surveillance Program. Antimicrob Agents Chemother, 2019. 63(7).

26. Mermel, L.A., et al., Guidelines for the Management of Intravascular Catheter-Related Infections. Clinical Infectious Diseases, 2001. 32(9): p. 1249–1272.

27. Xu, H., et al., Spectroscopy of single hemoglobin molecules by surface enhanced Raman scattering. Phys. Rev. Lett., 1999. 83: p. 4357.

28. Phillips, W.A., C.S. Hosking, and M.J. Shelton, Effect of ammonium chloride treatment on human polymorphonuclear leucocyte iodination. J Clin Pathol, 1983. 36(7): p. 808–10.

29. Lentacker, I., et al., Understanding ultrasound induced sonoporation: definitions and underlying mechanisms. Adv Drug Deliv Rev, 2014. 72: p. 49–64.

30. Wang, H.H., et al., Highly Raman-enhancing substrates based on silver nanoparticle arrays with tunable sub-10 nm gaps. Advanced Materials, 2006. 18(4): p. 491–495.

31. CLSI, Performance Standards for Antimicrobial Susceptibility Testing. 30th ed. CLSI document M100. 2020, Wayne, PA: Clinical and Laboratory Standards Institute.

32. Cushnie, T.P.T., N.H. O’Driscoll, and A.J. Lamb, Morphological and ultrastructural changes in bacterial cells as an indicator of antibacterial mechanism of action. Cellular and Molecular Life Sciences, 2016. 73(23): p. 4471–4492.

33. Mitchell, A.M. and T.J. Silhavy, Envelope stress responses: balancing damage repair and toxicity. Nature Reviews Microbiology, 2019. 17(7): p. 417–428.

34. Geisinger, E., et al., Acinetobacter baumannii: Envelope Determinants That Control Drug Resistance, Virulence, and Surface Variability. Annu Rev Microbiol, 2019. 73: p. 481–506.

35. Zgurskaya, H.I. and V.V. Rybenkov, Permeability barriers of Gram-negative pathogens. Ann N Y Acad Sci, 2020. 1459(1): p. 5–18.

36. Brauer, M.J., et al., Conservation of the metabolomic response to starvation across two divergent microbes. Proc Natl Acad Sci U S A, 2006. 103(51): p. 19302–7.

37. Link, H., et al., Real-time metabolome profiling of the metabolic switch between starvation and growth. Nature Methods, 2015. 12(11): p. 1091–1097.

38. Liebeke, M., et al., A metabolomics and proteomics study of the adaptation of Staphylococcus aureus to glucose starvation. Mol Biosyst, 2011. 7(4): p. 1241–53.

39. Drlica, K., et al., Quinolone-mediated bacterial death. Antimicrob Agents Chemother, 2008. 52(2): p. 385–92.

40. Snyder, M. and K. Drlica, DNA gyrase on the bacterial chromosome: DNA cleavage induced by oxolinic acid. J Mol Biol, 1979. 131(2): p. 287–302.

41. Khodursky, A.B., E.L. Zechiedrich, and N.R. Cozzarelli, Topoisomerase IV is a target of quinolones in Escherichia coli. Proc Natl Acad Sci U S A, 1995. 92(25): p. 11801–5.

42. Aldred, K.J., R.J. Kerns, and N. Osheroff, Mechanism of quinolone action and resistance. Biochemistry, 2014. 53(10): p. 1565–74.

43. Struve, C. and K.A. Krogfelt, Role of capsule in Klebsiella pneumoniae virulence: lack of correlation between in vitro and in vivo studies. FEMS Microbiology Letters, 2003. 218(1): p. 149–154.

44. Cantón, R., J.M. González-Alba, and J.C. Galán, CTX-M Enzymes: Origin and Diffusion. Frontiers in microbiology, 2012. 3: p. 110–110.

45. Spanu, T., et al., Evaluation of the new VITEK 2 extended-spectrum beta-lactamase (ESBL) test for rapid detection of ESBL production in Enterobacteriaceae isolates. Journal of clinical microbiology, 2006. 44(9): p. 3257–3262.

46. Yamaguchi, A., et al., Effects of magnesium and sodium ions on the outer membrane permeability of cephalosporins in Escherichia coli. FEBS Lett, 1986. 208(1): p. 43–7.

